# The impact of amplification on differential expression analyses by RNA-seq

**DOI:** 10.1101/035493

**Authors:** Swati Parekh, Christoph Ziegenhain, Beate Vieth, Wolfgang Enard, Ines Hellmann

## Abstract

Currently quantitative RNA-Seq methods are pushed to work with increasingly small starting amounts of RNA that require amplification. However, it is unclear how much noise or bias amplification introduces and how this effects precision and accuracy of RNA quantification. To assess the effects of amplification, reads that originated from the same RNA molecule (PCR-duplicates) need to be identified. Computationally, read duplicates are defined via their mapping position, which does not distinguish PCR- from natural duplicates and hence it is unclear how to treat duplicate reads.

Here, we generate and analyse RNA-seq datasets prepared using three different protocols (Smart-Seq, TruSeq and UMI-seq). We find that a large fraction of computationally identified read duplicates can be explained by sampling and fragmentation bias. Consequently, the computational removal of duplicates does not improve accuracy, power or FDR, but can actually worsen them. Even when duplicates are experimentally identified by unique molecular identifiers (UMIs), power and FDR are only mildly improved. However, the pooling of samples as made possible by the early barcoding of the UMI-protocol leads to an appreciable increase in the power to detect differentially expressed genes.

## Introduction

High throughput RNA sequencing methods (RNA-Seq) are currently under active development and might soon replace microarrays as the method of choice for gene expression quantification.^1–5^ For many applications, RNA-Seq technologies are required to become more and more sensitive; ideally, we would like to detect rare transcripts in single cells. Thereby, the sensitivity, accuracy and precision of transcript quantification strongly depends on how mRNA is converted into the cDNA that is eventually sequenced. In order to generate enough cDNA for sequencing, especially when starting from low amounts of RNA, amplification is often necessary.^6, 7^ Although it is known that PCR does not amplify all sequences equally well,^8–10^ PCR amplification is used in two popular RNA-Seq library preparation protocols (TruSeq & Smart-Seq^11^). However, it is still unclear how PCR bias in RNA-seq library preparation effects quantitative RNA-Seq analyses and to what extend PCR amplification adds noise and hence reduces the precision of transcript quantification. For detecting differentially expressed genes this is even more important than accuracy since it influences the power and potentially the false discovery rate.

RNA-Seq library preparation methods are designed with different goals in mind. TruSeq is the method of choice, if there is sufficient starting material, while the Smart-Seq protocol is better suited for low starting amounts.^12, 13^ Methods using UMIs and cellular barcodes have been optimized for low starting amounts and low costs to generate RNA-seq profiles from single cells.^6, 14^ To achieve these goals the methods differ in a number of steps, that will also impact the probability of read duplicates and their detection (Figure 1). TruSeq uses heat-fragmentation of mRNA and the only amplification is the amplification of the sequencing library. Thus all PCR duplicates can be identified by their mapping positions.

**Figure 1.**
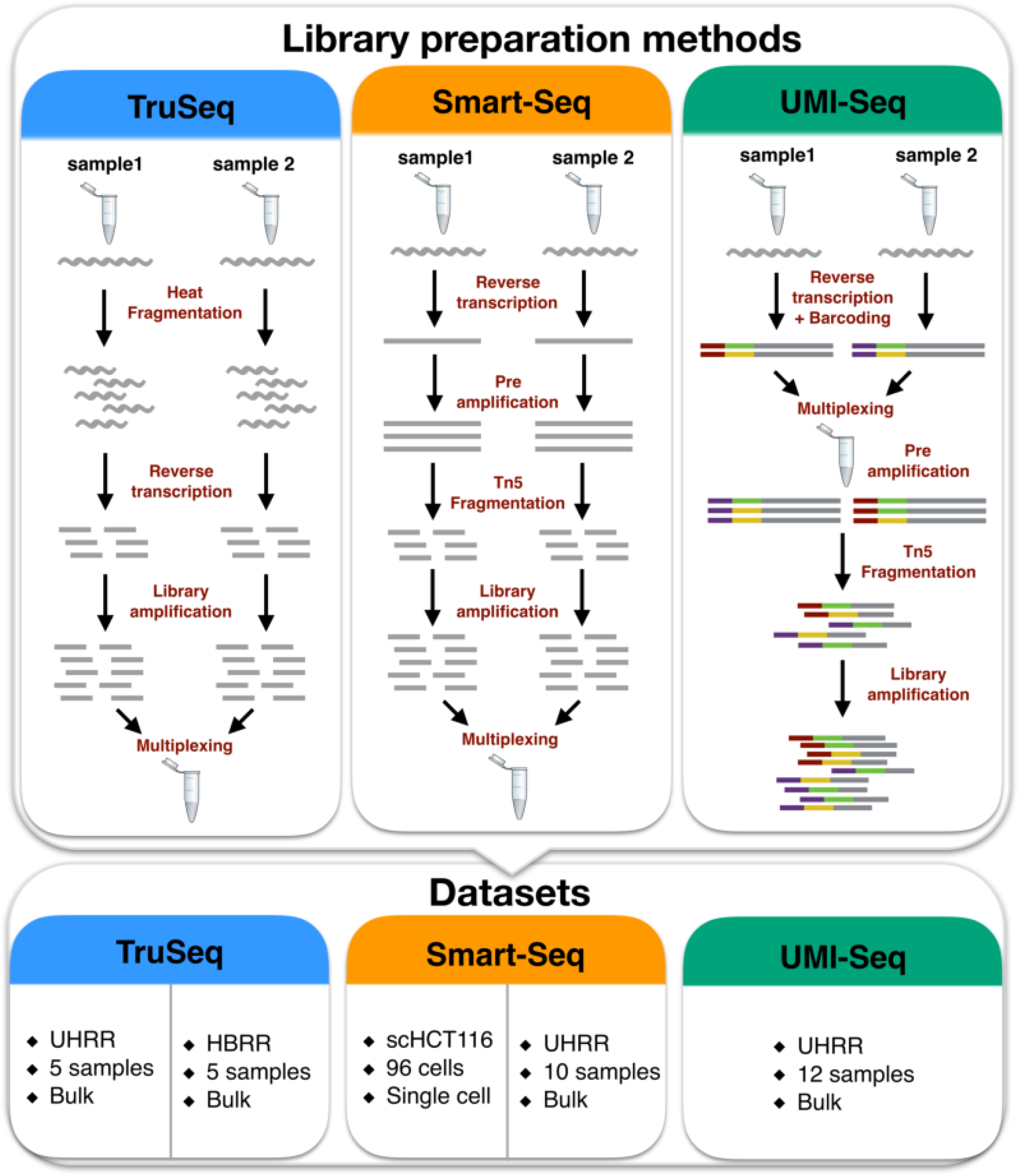
Schematic of library preparation protocols and datasets. The upper panel details the steps for the three different sequencing library preparation methods analysed in this study. In the UMI-Seq flow-chart red and purple tags represent the sample barcodes and the green and yellow tags the UMIs. In the lower panel, the datasets are described with the RNA-source and the number of replicates.

In the Smart-Seq protocol, full length mRNAs are reverse transcribed, pre-amplified and the amplified cDNA is then fragmented with a Tn5 transposase. Hence, PCR duplicates that arise during the pre-amplification step can not be identified by their mapping positions. UMI-Seq also generates cDNA by pre-amplification and Tn5 fragmentation,^11^ but unique molecular identifiers (UMIs) as well as library barcodes are already introduced during reverse transcription, i.e. before pre-amplification. This early barcoding allows all samples to be pooled after reverse transcription. The primer sequences required for the library amplification are introduced at the 3’end during reverse transcription. Thus, PCR-duplicates in UMI-seq data can always be identified via the UMI. Furthermore, computationally identified duplicates can also arise by sampling independent molecules. For a transcript of a given length, the chance for such “natural duplicates” increases with expression levels and fragmentation bias. In brief, for TruSeq-data duplicates can be identified computationally, while in Smart-Seq pre-amplification duplicates will escape detection and UMI-Seq is the only methods for which we know which reads are PCR duplicates.

That said, it is unclear whether removing duplicate reads computationally actually improves accuracy and precision by reducing PCR bias and noise or whether it decreases accuracy and precision by removing genuine information. Here, we investigate the impact of PCR amplification in RNA-seq by analyzing RNA-Seq datasets prepared with three different protocols (Smart-Seq, TruSeq and UMI-seq) and different amounts of amplification. We investigate the source of read duplicates by analysing PCR bias and fragmentation bias, assess accuracy by using ERCCs – spiked-in mRNAs of known concentrations^15^ and assess precision by power simulations using PROPER.^16^

## Results

### Selection of datasets

We analyse five different datasets, that represent three popular RNA-seq library preparation methods, starting with two benchmarking datasets from the literature.^2^ Both those datasets sequenced five replicates of bulk mRNA using the TruSeq protocol on commercially available reference mRNAs: the Universal Human Reference RNA (UHRR; Agilent Technologies) and the Human Brain Reference RNA (HBRR,ThermoFisher Scientific). We also used the UHRR samples to produce comparable Smart-Seq RNA-Seq and UMI-Seq data. All four bulk-RNA datasets high RNA-quality was used that also has comparable expression complexity and a low variance between the replicates (Table 1), thus ensuring good overall comparability of the datasets.

**Table 1.**
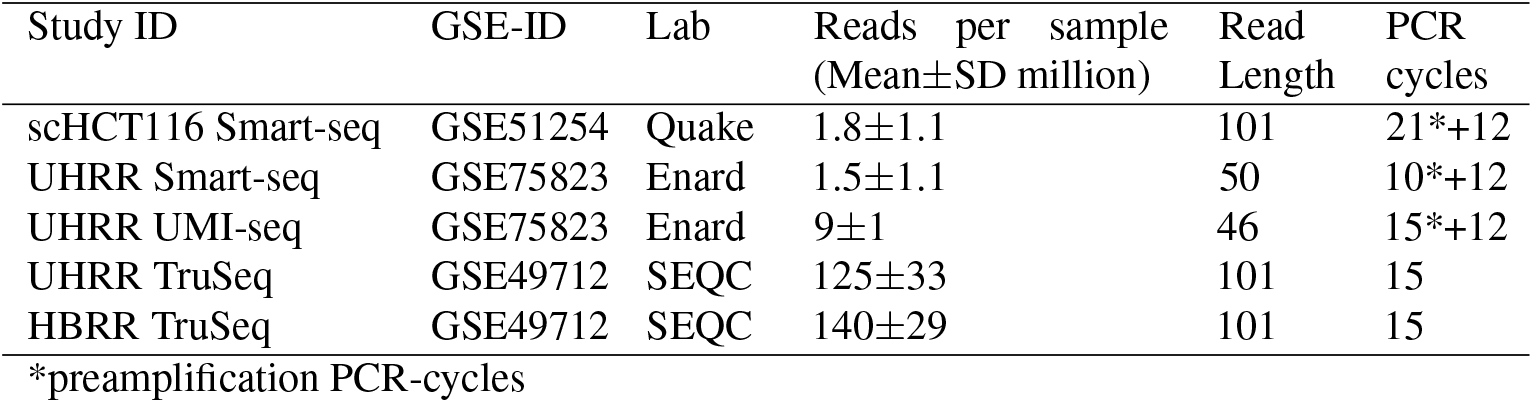
Description of the datasets analysed here.

| Study ID | GSE-ID | Lab | Reads per sample<br>(Mean±SD million) | Read Length | PCR cycles |
| --- | --- | --- | --- | --- | --- |
| scHCT116 Smart-seq | GSE51254 | Quake | 1.8±1.1 | 101 | 21*+12 |
| UHRR Smart-seq | GSE75823 | Enard | 1.5±1.1 | 50 | 10*+12 |
| UHRR UMI-seq | GSE75823 | Enard | 9±1 | 46 | 15*+12 |
| UHRR TruSeq | GSE49712 | SEQC | 125±33 | 101 | 15 |
| HBRR TruSeq | GSE49712 | SEQC | 140±29 | 101 | 15 |

\*preamplification PCR-cycles

Finally, we also wanted to include a single cell data and therefore chose the first published single cell dataset from Wu et al. (2014).^17^ The library preparation method used for the single cell data is also Smart-Seq and thus comparable to our UHRR-Smart-Seq data. However, the variance amongst replicates is expected to be higher, because those are biological and not technical replicates of a cancer cell line (HCT116).

All datasets contained ERCC-spike-ins, which allows us to compare the accuracy of the quantification of RNA-levels and all datasets except the UHRR-UMI-Seq have paired-end sequencing, which should provide more information for the computational identification of PCR duplicates.

### Natural duplicates are expected to be common

The number of computationally identified paired-end read duplicates (PE-duplicates) varies between 14% and 37% for the bulk data and 2% and 60% for the single cell data. Since single-end data is commonly used for gene expression quantification we also consider the mapping of the first read of every pair. The resulting fractions of computationally identified duplicates from single-end reads (SE-duplicates) are much higher: For the bulk data, it ranges from 40-76% and for the single cell data from 9-92% (Table 2, Figure 2a). For the UMI-Seq data, we make use of the design that each unique UMI sequence represents one RNA molecule of the original sample and find that UMI libraries show on average the highest duplicate fractions with 66% (Range:64-68%), whereas all those duplicates are bona-fide PCR-duplicates. In the UHRR Smart-Seq libraries, we only computationally identified 29% PE-duplicates (Figure 2a). Although these numbers are not strictly comparable due to differences in the library preparation (e.g. 5 more PCR-cycles for the UMI-data Table 1 and a stronger 3’ bias Supplementary Figure S4), it nevertheless strongly indicates that many PCR-duplicates in Smart-Seq libraries occur during pre-amplification and thus cannot be detected by computational means.

**Figure 2.**
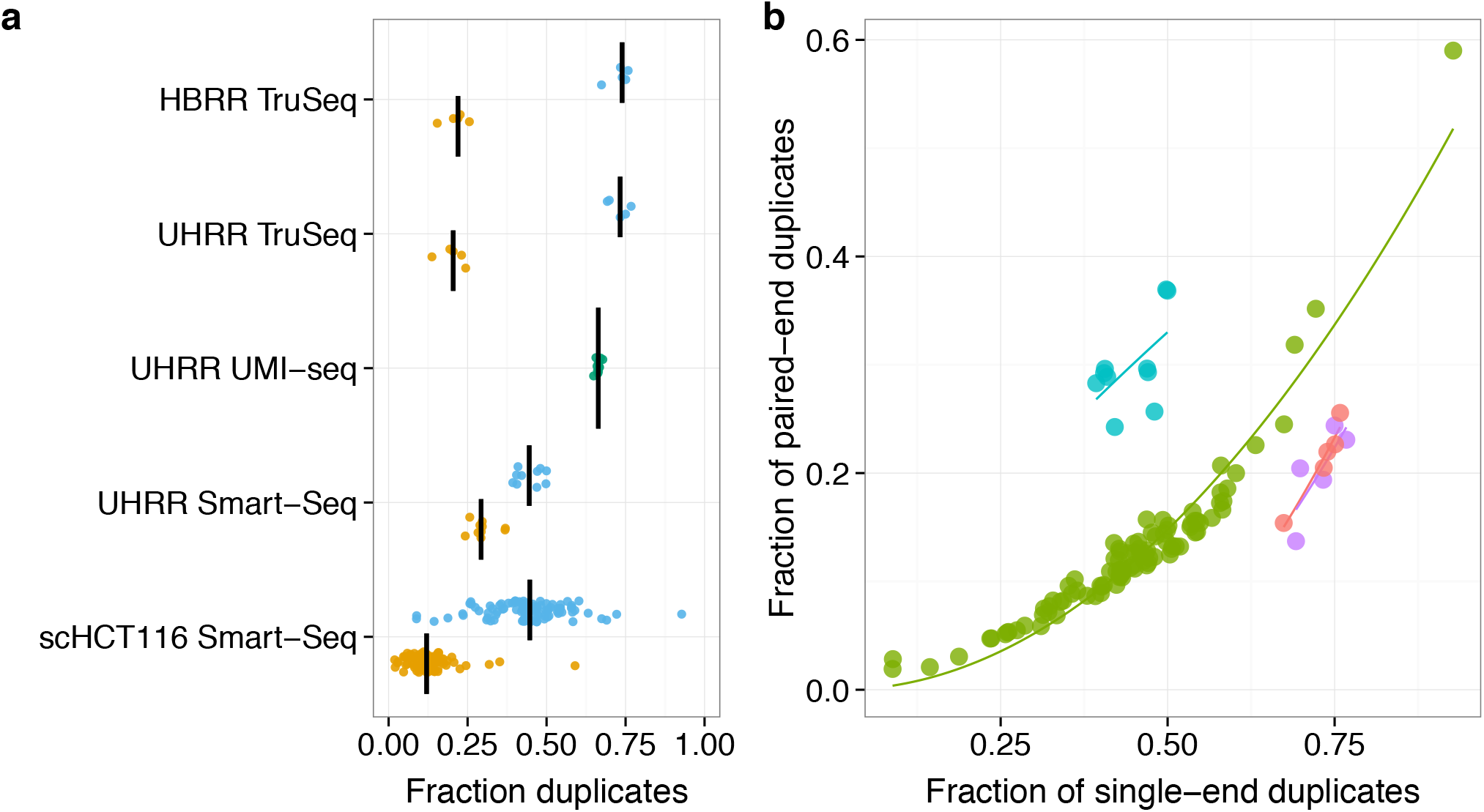
Fractions of SE- and PE- duplicates. In panel **a)**, we plot the fraction of computationally identified SE-duplicates (blue) and PE-duplicates (yellow) per sample. For the UMI-Seq data, we identify duplicates only based on the experimental evidence provided by the UMIs. The black line marks the median of the dataset. The relation between SE- and PE-duplicates is expected to follow a quadratic function, if the majority of duplicates are natural, i.e. due to fragmentation and sampling. In panel **b)**, we plot a quadratic fit for the different datasets (UHRR-TruSeq – purple, HBRR-TruSeq – red, UHRR-Smart-Seq – blue, scHCT116 – green).

**Table 2.** Fraction of duplicates per sample

| Study Name | Fraction PE-duplicates | Fraction SE-duplicates |
| --- | --- | --- |
| HBRR TruSeq | 0.15-0.26 | 0.67-0.76 |
| scHCT116 Smart-Seq | 0.02-0.59 | 0.09-0.92 |
| UHRR Smart-Seq | 0.24-0.37 | 0.40-0.50 |
| UHRR TruSeq | 0.14-0.24 | 0.70-0.77 |
| UHRR UMI-seq | 0.65-0.68* |  |

\* Fraction of duplicates based on UMI counts.

Generally, the fraction of duplicate reads is expected to depend on library complexity, fragmentation method and sequencing depth. Sequencing depth is the factor that gives us the most straight-forward predictions and in the case of SE-duplicates they are by and large independent of other parameters such as the fragment size distribution. As expected, we observe a positive correlation between the number of reads that were sequenced and the fraction of SE-duplicates (Figure 3). In order to test to what extend simple sampling can explain the number of SE-duplicates, we calculate the expected fraction of SE-duplicates, given the observed number of reads per gene and the gene lengths (see Methods, Figure 3). Note that, in the case of Smart-Seq this approach will only evaluates the effect of the library PCR, but be oblivious to PCR duplicates that arose during pre-amplification. We find that for TruSeq and Smart-seq the majority of SE-duplicates are expected under this simple model of random sampling (Figure 3). For the TruSeq data we underestimate the fraction of duplicates on average by 10% (8-13%), for the single cell Smart-Seq data by 11% (0.4-53%) and for the bulk Smart-Seq data by 18% (14-24%). Thus, irrespective of the library preparation protocol, a large fraction of computationally identified SE-duplicates could easily be natural duplicates (Figure 3).

**Figure 3.**
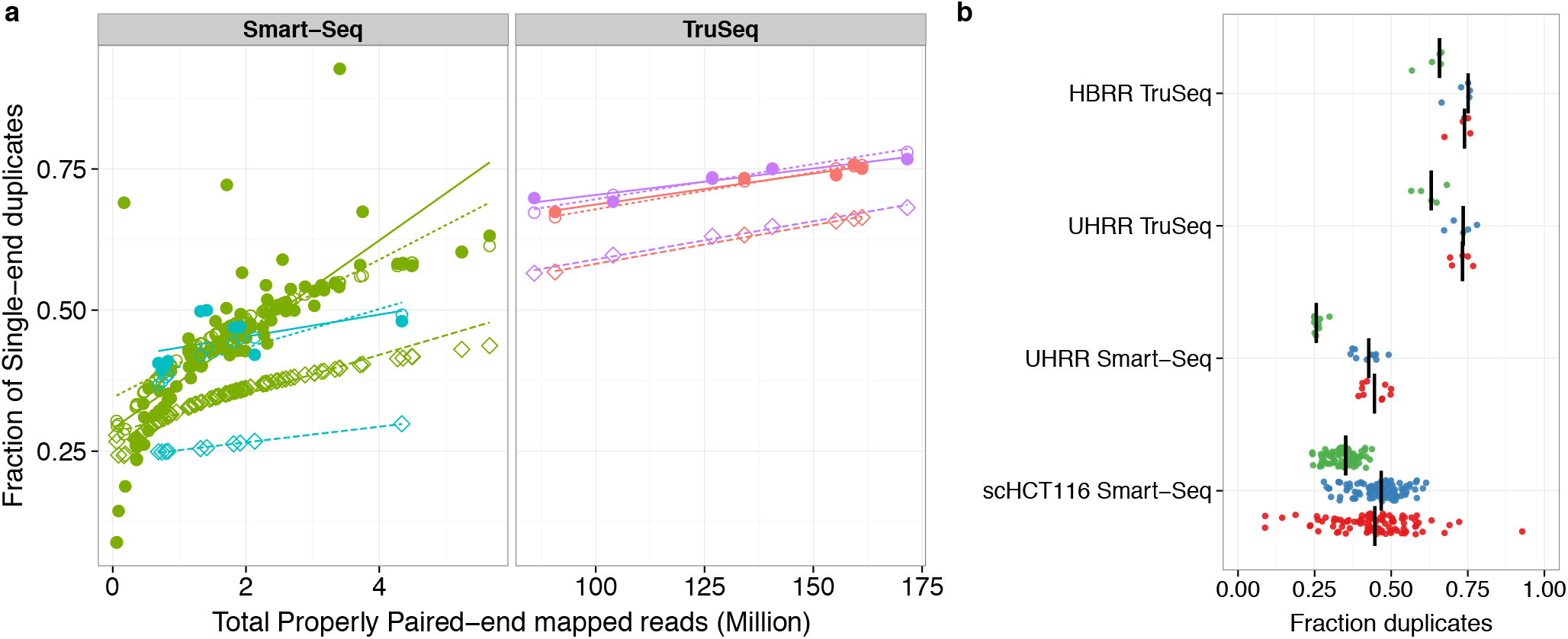
The Fraction of SE-duplicates increases with the total number of reads. If the correlation between sequencing depth and duplicates is due to sampling and fragmentation, we can thus quantify the impact of sampling and fragmentation. **a**) The left panel shows the two Smart-Seq datasets (UHRR- blue, scHCT116- green) and the right panel the TruSeq data (HBRR- red, UHRR- purple). Filled circles represent the observed fraction of SE-duplicates. Open symbols represent simulated data: Open diamonds mark the expected fractions of SE-duplicates under a simple sampling model and open circles are the expectations for a sampling model with fragmentation bias. The lines are the log-linear fits between sampling depth and SE-duplicates per dataset. In **b**), we plot only the SE-duplicate fractions (as in a) for an easier comparison between the observed (red) and expected (sampling – green, sampling+fragmentation – blue).

In contrast to this simple sampling expectation for SE-duplicates, fragments produced during PCR-amplification after adapter ligation, will necessarily produce fragments with the same 5’ and 3’ end and consequently will have identical mapping for both ends. If the sampling was shallow enough so that we would not expect to draw the same 5’ end twice by chance, the 3’end position should also be identical and no reads with only one matching 5’end are expected. If same 5’ ends are more frequent due to biased fragmentation, we expect a higher ratio of SE- to PE-duplicates. Thus, the relationship between PE- and SE-duplicates contains information about the relative amounts of duplicates produced by fragmentation as compared to amplification. More specifically, we expect that the fragmentation component of the PE- vs. SE-duplicates should be captured by a quadratic fit with an intercept of zero.

The quadratic term is not significant for the UHRR-Smart-Seq and the UHRR-TruSeq data, which could be due to shallow sequencing and low sample size, but could also be seen as an indication of a higher proportion of PCR-duplicates. On the other hand, the quadratic term is significant and positive for the HBRR TruSeq and the scHCT116 datasets, supporting the notion that at least for those datasets library PCR amplification is not the dominant source of duplicates. This is also consistent with our finding that most observed SE- duplicates are simply due to sampling (Supplementary Table S1 and Figure 2).

### Fragmentation is biased

If fragmentation does not occur randomly but some sites are more likely to break than others, those might increase the fraction of SE-duplicates. To evaluate the impact and nature of fragmentation bias, we analysed ERCC spike-ins because they are exactly the same in all datasets. First, we test whether the variance in the frequency of 5’ end mapping positions of ERCCs in one sample can explain a significant part of this variance in other samples prepared with the same method. We find on average an of 0.77 and 0.85 for the Smart-Seq and TruSeq protocols, respectively. Note, that this high *R*^2^ holds for samples that were prepared in different labs, for example the *R*^2^ between the Smart-Seq samples prepared in our lab and the single cell data from the Quake lab ranges between 0.56-0.90. In contrast, if the *R*^2^ is calculated for the comparison between one TruSeq and one Smart-Seq library, it drops to 0.0012 (Figure 4 a,b). All in all, this is strong evidence that fragmentation prefers reproducibly the same sites given a library preparation protocol and thus read sampling is not random.

**Figure 4.**
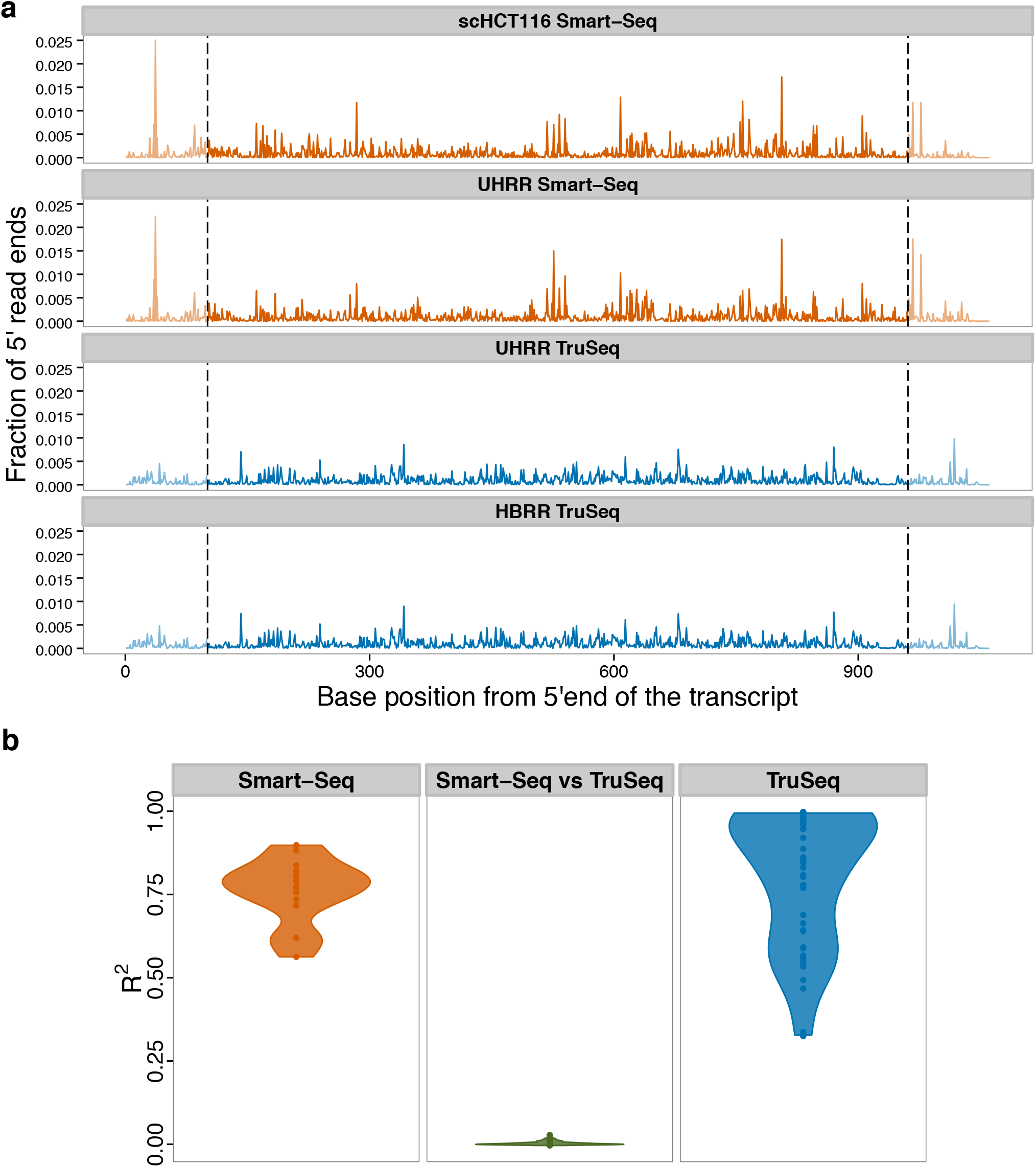
The fragmentation patterns of the ERCCs are highly reproducible for different samples prepared with the same RNA-seq library method. **a**) Here, we plot the fraction of 5’read ends per position of ERCC-00002. Because the TruSeq libraries (blue) had read lengths of 100 bases, we do not consider the ends (grey dashed lines) for the calculation of the pair-wise *R*^2^ values for fragmentation. **b**) Violin plot of the adjusted *R*^2^ of a linear model of 5’read ends from different samples. The reproducibility of fragmentation is highest between Smart-Seq samples (orange), a little lower between the TruSeq samples and there is no correlation between samples from one Smart-Seq and one TruSeq sample (middle,green).

To identify potential causes for these non-random fragmentation patterns, we correlated the GC-content of the 15 bases around a given position with the number of 5’ read ends. This explained very little of the fragmentation patterns in the TruSeq-data (median *R*^2^ = 0.0064, 59% of the pair-wise comparisons significant with *p* < 0.05), and none in the Smart-Seq data (median *R*^2^ = 0.00002, 18% significant with *p* < 0.05, Supplementary Figure S1a and Supplementary Table S2). Next, we built a binding motif for the Transposase^18^ from our UHRR-Smart-Seq data and, unsurprisingly, found that the motif has a very low information content (Supplementary Figure S1b) and accordingly a weak effect on the 5’ read end count (median *R*^2^ = 0.0019, 48% & 58% significant with *p* < 0.05 for scHCT116 & UHRR Smart-Seq, Supplementary Figure S1a and Supplementary Table S2).

Although, we could not identify the cause for the fragmentation bias in the sequence patterns around the fragmentation site, we can still quantify the maximal impact of fragmentation bias on the number of SE-duplicates, simply by adjusting the effective length of the transcripts. For the TruSeq data, we estimate that a fragmentation bias that reduces the effective length approximately ∽ 2-fold gives a reasonably good fit, leaving on average 1% ( 0.1-2.6%) of the SE-duplicates unexplained. For the UHRR-Smart-Seq data, a ∽ 36-fold reduction in the effective length is needed and leaves only 3% (1.1-7.7%) of the duplicates unexplained. For the single cell data, the fragmentation bias that gives overall the best fit is a ∽6-fold reduction, however the fit is worse since the fraction of unexplained duplicates is still at ∽5% and varies between 1% and 41% (Figure 3). In summary, we find that fragmentation bias contributes considerably to computationally identified read duplicates and is stronger for Smart-Seq, i.e. for enzymatic fragmentation, than for TruSeq, i.e. heat fragmentation.

### Removal of duplicates does not improve the accuracy of quantification

To evaluate the impact of PCR duplicates on the accuracy of transcript quantification, we use again the ERCC spike-in mRNAs: Although, the absolute amounts of ERCC-spike ins might vary due to handling, the relative abundances of these 92 reference mRNAs can serve as a standard for quantification. Ideally, the known concentrations of the ERCCs should explain the complete variance in read counts and any deviations are a sign of measurement errors. We calculate the *R*^2^ values of a log-linear fit of transcripts per million (TPM) versus ERCC concentration to quantify how well TPM estimates molecular concentrations and compare the fit among the different duplicate treatments. In no instance does removing read duplicates improve the fit, but in most cases it gets significantly worse (t-test, *p* < 2 x 10^-3^) except for for the computational PE-duplicate removal of the UHRR-Smart-Seq and the duplicate removal using UMIs (Figure 5a).

### Removal of duplicates does not improve power

Most of the time we are not interested in absolute quantification, but are content to find relative differences, i.e. differentially expressed (DE) genes between groups of samples. The extra noise from the PCR-amplification has the potential to create false positives as well as to obscure truly DE genes. In order to assess the impact of duplicates on the power and the false discovery rate to detect DE genes, we simulate data based on the estimated gene expression distributions of the five datasets. For comparability, we first equalized the sampling depth by reducing the number of mapped reads to 3 million and 1 million for bulk and single cell data, respectively. Next, we estimated gene-wise base mean expression and dispersion using DESeq2.^19^

There are no big differences in the distributions of mean baseline expression and dispersion estimates from the different duplicate treatments for the two Smart-Seq datasets, whereas there is a shift towards lower means and higher dispersion, when removing SE-duplicates for the TruSeq datasets. Dispersions shift only to lower values if we exclude duplicates based on identification by UMIs (Figure 6a, Supplementary Figure S2). The empirical mean and dispersion distributions are then used to simulate two groups with six replicates for bulk-RNA-Seq datasets and 45 replicates for the single cell dataset. In all cases we simulate that 5% of the genes are differentially expressed with log2-fold changes drawn from a normal distribution with N(0,1.5).^16^ We analysed 100 simulations per data-set using DESeq2 and calculate the power and the FDR for detecting DE-genes with a log2-fold change of at least 0.5.

**Figure 6.**
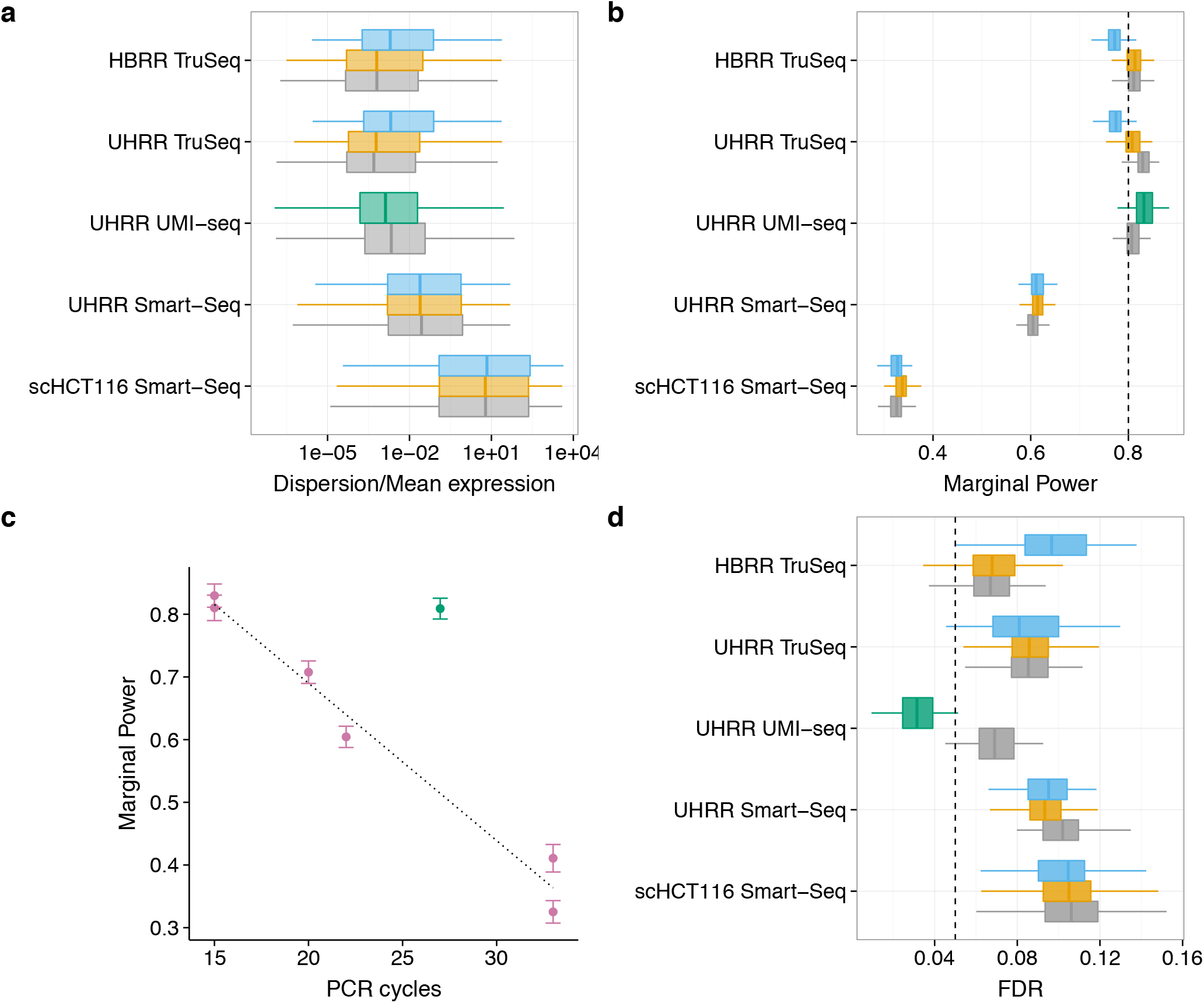
Duplicate removal has little influence on the power and FDR to detect DE-genes in comparison to the library preparation method. We estimated the distributions of mean expression and dispersion across genes for each dataset using DESeq2 after downsampling the datasets to 3 or 1 million reads. The distributions are estimated for the data including all reads (grey), removing PE-duplicates (yellow), removing SE-duplicates (blue) and for the UHRR-UMI-Seq dataset removing duplicates using UMIs (green). In analogy to the coefficient of variation, we summarize distributions of dispersion/Mean in the boxplot **a**. The estimated mean and dispersion distributions served as input for our power simulations using PROPER.^16^ We did 100 simulations per dataset, whereas each dataset had two groups of six replicates (45 for scHT116) with 5% of the genes being differentially expressed between the groups. In panel **b**, we report the marginal power to detect a log2-fold change of 0.5 and in panel **d** the corresponding FDR, whereas the nominal FDR was set to *α* = 0.05 (dashed line). In panel **c**, we plot our estimates of the Marginal power against the number of PCR-cycles for each dataset. Error bars are standard deviation to the mean marginal power over 100 simulations. We find a surprisingly simple linear decline in power with the number of PCR-cycles, if we only consider datasets where PCR amplification was done separately for each sample of the dataset (violet). To confirm this simple fit we added two other datasets: 1) Bulk Smart-Seq dataset of mouse brain bulk RNA amplified using 20 PCR-cycles and 2) Single cell Smart-Seq dataset of 96 mouse embryonic stem cells that were amplified using 33 cycles. The only outlier is the UMI-Seq dataset for which samples were pooled prior to amplification (green).

Except for the UHRR-UMI-seq dataset, the nominal FDR that we set to *α* = 5% is exceeded: it varies between 7.5% and 13.3 %, whereas the HBRR TruSeq has the lowest and the scHCT116 Smart-Seq data has the highest FDR (Figure 6d). Computational removal of the duplicates has very little impact on the FDR, which is significant for only two datasets: In the HBRR-TruSeq dataset SE-duplicate removal increases the FDR by 3% and for the UHRR-Smart-Seq data PE- and SE-duplicate removal improve the fit by 1% (Figure 6d). Again, the only convincing improvement is achieved by duplicate removal using UMIs, which reduces the FDR from 7% to 3%. (t-test, *p* < 1 x 10^-15^).

The differences in the power are more striking. As for the FDR, the major differences are not between duplicate treatments, but between the datasets. For the TruSeq and the UHRR-UMI datasets, the average power to detect a log2-fold change of 0.5 is ∽ 80% (Figure 6b). For those datasets the changes in power due to duplicate removal are only marginal and for the computational removal using PE-duplicates it actually decreases the power for the UHRR-TruSeq datasets by 2%, while for the UMI-seq data duplicate removal increases power by 2%. The power for the UHRR-Smart-Seq and the scHCT116 Smart-Seq datasets is with 60% and 33%, respectively, much lower, and duplicate removal increases the power by only 1%. The large differences in power between the datasets are unlikely to be ameliorated by increasing the number of replicates per group: Additionally to the 6 and 45 replicates for which the results are reported above, we also conducted simulations for 12 and 90 replicates for bulk and the single cell data, respectively. This doubling in replicate number increases the power for the UHRR-Smart-Seq dataset only from 60 to 67% and for the single cell dataset from 33 to 35% (Supplementary Figure S5, Supplementary Table 3).

## Discussion

RNA-Seq has become a standard method for expression quantification and in most cases, the sequencing library preparation involves amplification steps. Ideally, we would like to count the number of RNA molecules in the sample and thus would want to keep only one read per molecule. A common strategy applied for amplification correction in SNP-calling and ChIP-Seq protocols^20, 21^ is to simply remove reads based on their 5’ends, so called read duplicates. Here, we show that this strategy is not suitable for RNA-Seq data, because the majority of such SE-duplicates is likely due to sampling. For highly transcribed genes, it is simply unavoidable that multiple reads have the same 5’end, also if they originated from different RNA-molecules. We find that only 10% and max. 20% of the read duplicates cannot be explained by a simple sampling model with random fragmentation. This fraction decreases even more, if we factor in our finding that the fragmentation of mRNA or cDNA during library preparation is clearly non-random: We find a strong correlation between the 5’ read positions of the ERCC-spike-ins across samples. Because local sequence content has little or no detectable effect on fragmentation, we cannot predict fragmentation, but we can quantify the observed effect: a fragmentation bias that halfs the number of break points can fit the observed proportion of duplicates for TruSeq libraries well. For the Smart-Seq datasets, fragmentation biases would have to be much higher to explain the observed duplicates, fit the observed duplicate fractions less well and are also inconsistent between the datasets (35 for the UHRR and 6 for the scHCT116).

Since computational methods cannot distinguish between fragmentation and PCR duplicates, the removal of duplicates could introduce a bias rather than removing it. Using the ERCC-spike-ins, we can indeed show that removing duplicates computationally does not improve a fit to the known concentrations, but rather makes it worse, especially if only single-end reads are available (Figure 5). This is in line with our observation that most single end duplicates are due to sampling and fragmentation, and removing duplicates is thus equivalent to a saturation effect known for microarrays.

**Figure 5.**
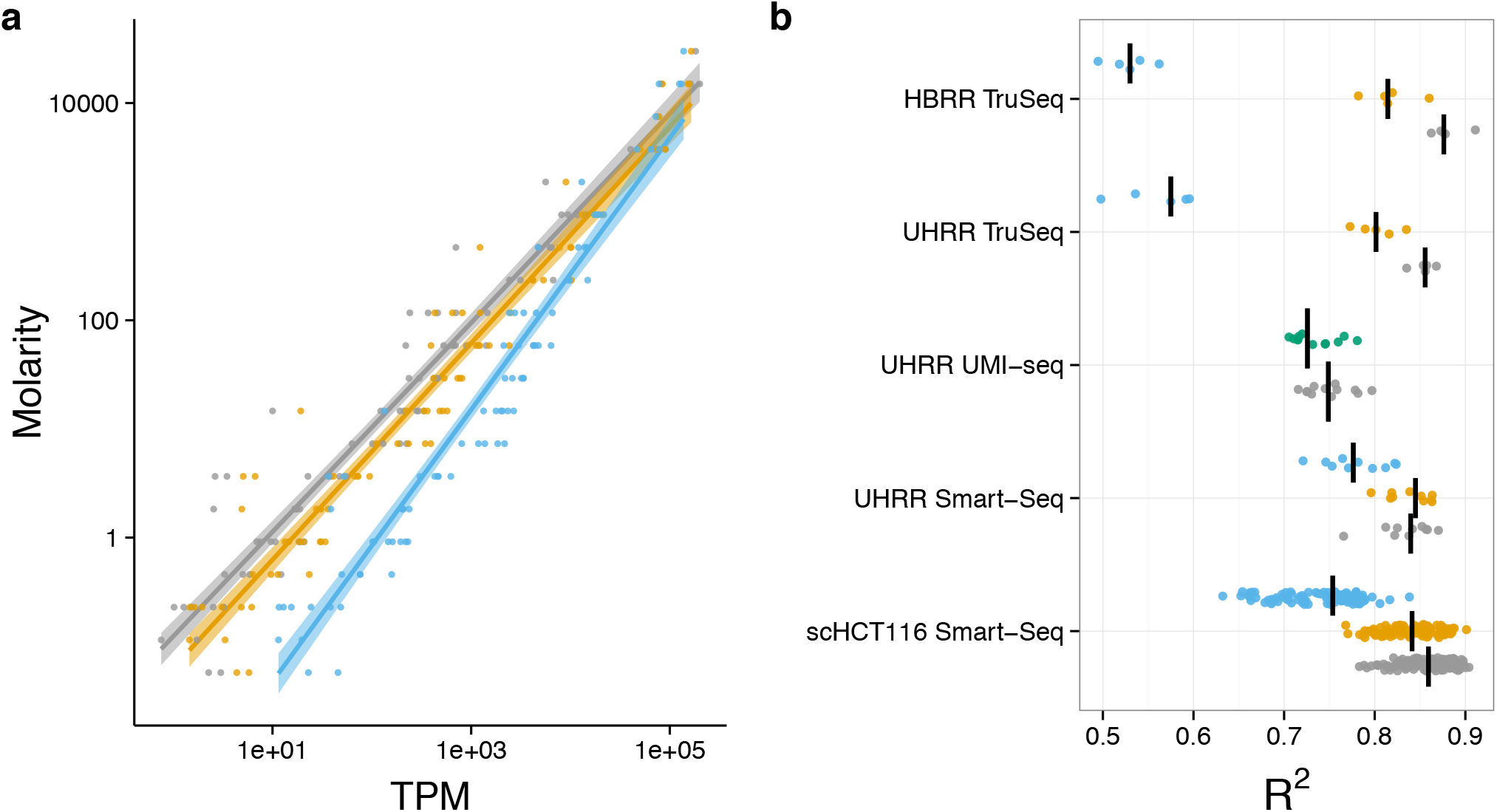
Removing duplicates does not improve the accuracy of expression quantification as measured using the ERCC spike-ins. Expression levels as quantified in transcripts per million reads (TPM) are a good predictor of the concentrations of the ERCC spike-ins. The log-linear fit of TPM vs. Molarity for one exemplary sample of the UHRR-TruSeq dataset is shown in **a)**. The most accurate prediction of ERCC molarity is the TPM estimator using all reads (grey). Removing duplicates as PE (yellow) makes the fit a little worse and removing SE-duplicates (blue) much worse. The adjusted *R*^2^ for all samples are summarized in **b)**, the median for each dataset is marked as black line. The *R*^2^ of the TPM estimate from the removal of PCR-duplicates using UMIs (green) is surprisingly similar to keeping PCR-duplicates (grey).

Moreover, the Smart-Seq protocol, which was designed for small starting amounts, involves PCR amplification before the final fragmentation of the sequencing library. Thus in the case of Smart-Seq, computational methods cannot identify PCR duplicates that occur during this pre-amplification step. When we use unique molecular identifiers (UMIs), we find that 66% of the reads are PCR duplicates and only 34% originate from independent mRNA molecules. In contrast, when using paired-end mapping for a comparable Smart-Seq library, we identify 27% as duplicates and 73% as unique. Since mainly 3’ends of transcripts are sequenced in UMI-seq, the complexity of the library is probably decreased, which increases the fraction of PCR duplicates for a given sequencing depth. However, it seems unlikely that library complexity can explain the 30% difference in duplicate occurrence. Rather it seems likely that the difference is due to PCR-duplicates that are generated during pre-amplification and thus remain undetectable by computational means.

In summary, computational methods are limited when it comes to removing PCR-duplicates, but how much noise or bias do PCR duplicates actually introduce? In other words, the metric that we are actually interested in is how PCR-duplicates impact the power and the false discovery rate for the detection of differentially expressed genes. Both, power and FDR, are determined by the gene-wise mean expression and dispersion. Based on simulated differential expression using the empirically determined mean and dispersion distributions, we find that computational removal of duplicates has either a negligible or a negative impact on FDR and power, and we therefore recommend not to remove duplicates. If PCR duplicates are removed using UMIs, both FDR and power improve, although the effects in the bulk data analysed here are relatively small: FDR is improved by 4% and the power by 2%. However, it is likely that UMIs get more important when using small amounts of starting material, as it is the case for single-cell RNA-seq.^22^

The major differences in power are between the datasets, with the TruSeq data and the UMI data giving 20% and 40% higher power than the Bulk and the single cell Smart-Seq data, respectively. One possible explanation for the differences in power is the total number of PCR-cycles involved in the library preparation. With every PCR-cycle the power to detect a log2-fold change of 0.5 appears to drop by 2.5% (Figure 6c). The only exception is the UMI-Seq dataset, that even if duplicates are not removed gives a power of 80%, which is comparable to the power reached with TruSeq despite the UMI-Seq method having 12 more PCR-cycles. Technically, UMI-Seq is most similar to the Smart-Seq method. The biggest difference between the two is that all UMI-Seq libraries are pooled before PCR-amplification, suggesting that the PCR-noise is due to the different PCR-reactions and not due to amplification efficiency per-se.

We conclude that computational removal of duplicates is not recommendable for differential expression analysis and if sufficient starting material is available so that only few PCR-cycles are necessary, the loss in power due to PCR duplicates is negligible. However, if more amplification is needed, power would be improved if all samples are pooled early on, and for really low amounts as for single cell data also the gain in power that is achieved by removing PCR-duplicates using UMIs will become important.

## Methods

### Datasets

We used six datasets representing the TruSeq, Smart-Seq and UMI-seq protocols and varying amounts of starting material from bulk RNA or single cell RNA. All analysed datasets contain the ERCCs spike-in RNAs. This is a set of 92 artifical poly-adenylated RNAs designed to match the characteristics of naturally occurring RNAs with respect to their length (273-2022bp), their GC-content (31-53%) and concentrations of the ERCCs (0.01-30,000 attomol/μl). The recommended ERCC spike-in amounts give 5 – 10^7^ ERCC RNA molecules in the cDNA synthesis reaction.

To reduce biological variation, we used the well-characterized Universal Human Reference RNA (UHRR; Agilent Technologies) for the two datasets produced for this study. We downloaded UHRR- and HBRR-TruSeq data from SEQC/MAQC-III.^2^ Finally, we also analyse the single cell data published in Wu et al. 2014,^17^ for which the colorectal cancer cell-line HCT116 was used (Table 1). The input mostly being commercially distributed human samples, we expect all biological samples analysed in this study to have similarly high quality and complexity. All data that were generated for this project were submitted to GEO under accession GSE75823.

### RNA-seq library preparation and sequencing

For the 10 Smart-Seq libraries, 250 ng of Universal Human Reference RNA (UHRR; Agilent Technologies) and ERCC spike-in control mix I (Life Technologies) were used and cDNA was synthesized as described in Picelli et al,^12^ but here we only use 9 PCR cycles for amplification. 1 ng of pre-amplified cDNA was used as input for Tn5 transposon tagmentation by the Nextera XT Kit (Illumina), followed by 12 PCR cycles of library amplification. For sequencing, equal amounts of all libraries were pooled.

For the UMI-Seq libraries, we started with 10 ng of UHRR-RNA to synthesise cDNA as described in Soumillon et al.^23^ This protocol is very similar to the Smart-Seq protocol, however the first strand cDNA is decorated with sample-specific barcodes and unique molecular identifiers. Barcoded cDNA from all samples was pooled, purified and unincorporated primers digested with Exonuclease I (NEB). Pre-amplification was performed by single-primer PCR for 15 cycles. 1 ng of full-length cDNA was then used as input to the Nextera XT library preparation modified to enrich for barcoded 3’ ends by addition of a custom i5 primer.

Library pools were sequenced on a Illumina HiSeq1500. For the Smart-Seq libraries were sequenced using 50 cycles of paired-end sequencing on a High-Output flow-cell. The UMI-seq libraries were sequenced on a rapid flow-cell, where the first read contains the sequences of the sample barcode and the UMI using 17 cycles. The second read sequence is the actual cDNA fragment with 46 cycles.

### Data Processing

For Smart-Seq and TruSeq libraries, the sequenced reads were mapped to the human genome (hg19) and ERCC reference sequences using NextGenMap^24^ by local alignment using the default parameters, except the following three the maximum fragment size which was set to 10kb, the minimum identity set to 90% and only the best hit per read was reported. The mapped reads were assigned to genes [Ensembl database annotation version GRCh37.74] using FeatureCount from the bioconductor package Rsubread.^25^

For UMI-seq libraries, cDNA reads were mapped to the Ensembl transcriptome [version GRCh37.74] also using NextGenMap. If either the sample barcode or the UMI had at least one base with sequence quality ≤ 10 or contained ‘N’s the read was discarded. Next, we generated count tables for Ensembl genes, using read counts or counts of unique UMI-sequences per gene.

Finally, mitochondrial and ambiguously assigned reads were remove from all libraries.

### Duplicates detection and removal

To flag single-end duplicates (SE), we used in-house scripting to identify reads that map to the same 5’ position, have the same strand and the same CIGAR value. Because we cannot determine the exact mapping position for 5’ soft clipped reads, we discard them. To flag paired-end duplicates (PE), we used the same requirements as for the SE-duplicates, those requirements had just to be fulfilled for both reads of a pair.

### Model for the fraction of sampling and fragmentation duplicates

We obtain an expectation for the number of reads if duplicates are identified via their 5’position and only one read per 5’end position is kept. We use the observed numbers of reads per gene (*r_G_*) and the effective length of the gene (*L_eG_* = *L* – 2 × readlength). Then the expected number of unique reads is

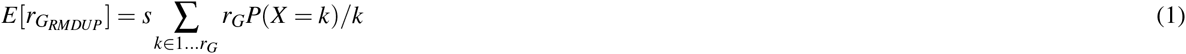

whereas *P*(*X* = *k*) derived from a positive Poisson distribution with *λ_G_* = *r_G_/L_eG_* and s is a scaling factor *s* = 1/Σ_*k*∊1…*rG*_ *P*(*X* = *k*).

In order to estimate the level of fragmentation bias, we simply modified the effective length *L_eG_* by a factor *f × L_eG_*.

### Fragmentation pattern analysis

To compare fragmentation sites across libraries, we counted 5’ read starts per position for the ERCCs across all datasets using samtools and in house perl scripts. To avoid edge effects in later analyses, we excluded the first and last 100 bases of each ERCC, whereas 100 bases is the maximum read length of datasets analysed here.

We generated a Position Weight Matrix (PWM) for the transposase (Tn5) motif by simply stacking up the 30 bases of the putative Transposase binding sites from all UHRR-Smart-Seq reads. Those 30 bases are identified as 6 bases upstream of the 5’ read end and the 24 downstream.^18^ The resulting PWM was then used to calculate motif scores across the ERCCs using the Bioconductor package PWMEnrich^26^.

### Power evaluation for differential expression

For power analysis, we estimated the mean baseline expression and dispersion for all datasets after downsampling them to 3 and 1 million reads for bulk and single cell data, respectively. This was done for all three duplicate treatments (keep all, remove SE and remove PE) using DESeq2^19^ with standard parameters. Furthermore, genes with very low dispersions (< 0.001) were removed. We chose the sample sizes 3, 6 and 12 per condition for the bulk data and 30, 45 and 90 for the single cell dataset, because they seemed to be a good representation of the current literature. For the simulations, we use an in-house adaptation of the Bioconductor-package PROPER.^16^ As suggested in Wu et al. 2015^16^, we set the fraction of deferentially expressed genes between groups to 0.05 and the log2-fold change for the DE-genes was drawn from a normal distribution with *N*(0,1.5). We generated 100 simulations per original input data-set and analysed them using DESeq2. Next, we calculated the power to detect a log2-fold change of at least 0.5 and the according FDR using *α* = 0.05.

## Acknowledgements

We thank Khalis Afnan and Sabrina Weser for help with the RNA-Seq library preparation.

## Author contributions statement

SP and CZ conceived the study. CZ prepared RNA-Seq libraries. SP, IH and BV analysed the data. IH, SP and WE wrote the manuscript. All authors read and approved the final manuscript.

## Additional information

### Accession codes

RNA-seq data generated for this study is submitted to GEO under the accession code: GSE75823.

The impact of amplification on differential expression analyses by RNA-seq

Swati Parekh, Christoph Ziegenhain, Beate Vieth, Wolfgang Enard, Ines Hellmann*

Anthropology & Human Genomics, Department of Biology II, Ludwig-Maximilians University, Großhaderner Str. 2, 82152 Martinsried, Germany.

*

## Supplementary figures

**Figure S1:**
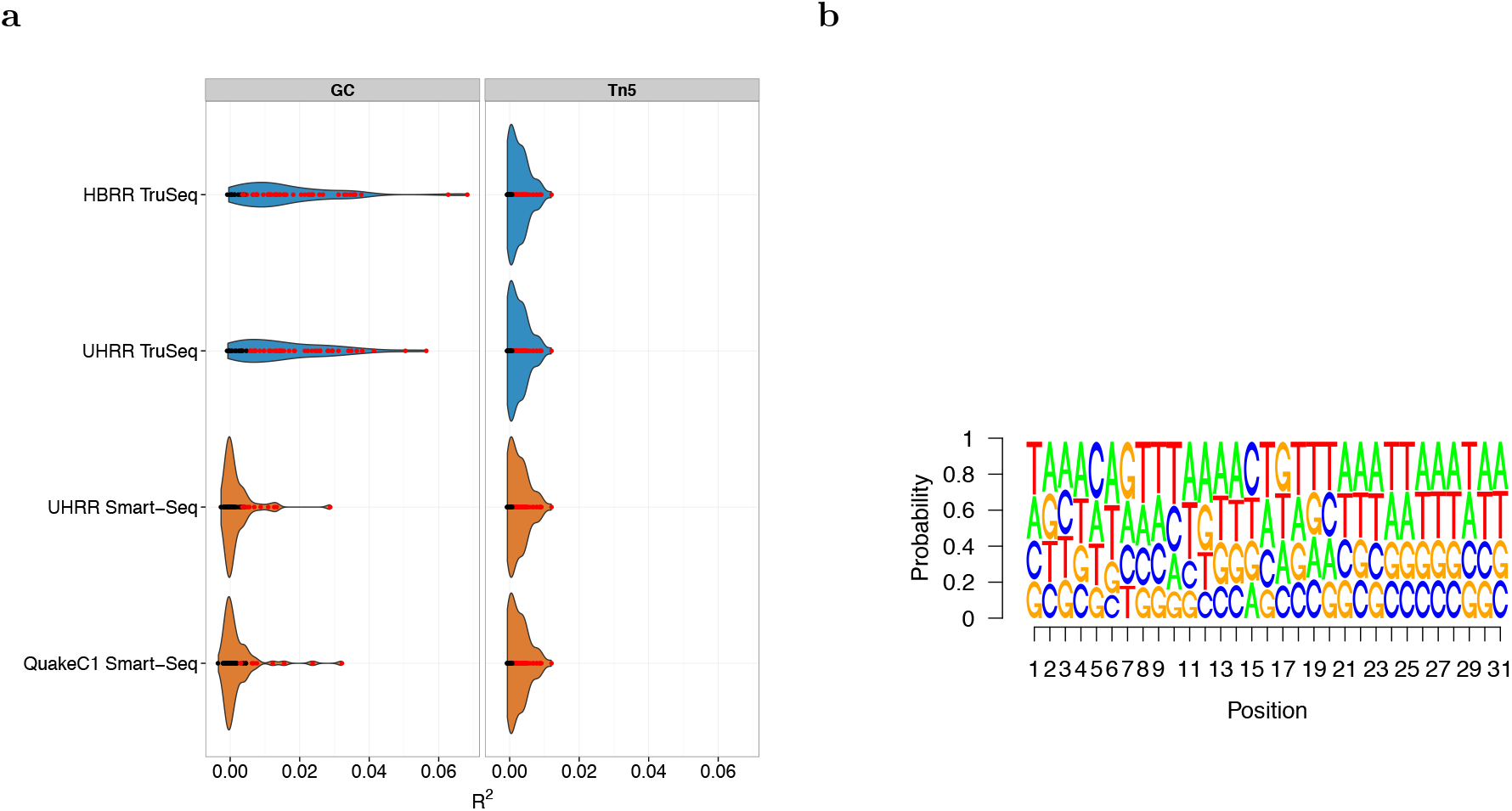
Fragmentation does not appear to have a cutting site preference. Colors of the violin plots represent library preparation methods, ‘blue – Smart-Seq, ‘orange’ – TruSeq and dots are colored by the significance of the fit where ‘red’ – pvalue ≤ 0.05 and ‘black’ – pvalue > 0.05. a) Left panel shows the violin plot of adjusted *R*^2^ of linear model fit between background corrected GC content and 5’ mapped read depth of the middle base in the 15bases window and on the right panel the adjusted *R*^2^ of linear model fit between the Tn5 motif score and 5’ mapped read depth calculated for ERCC spike-in RNAs. b) Sequence logo of the Tn5 motif derived from UHRR Smart-Seq dataset.

**Figure S2:**
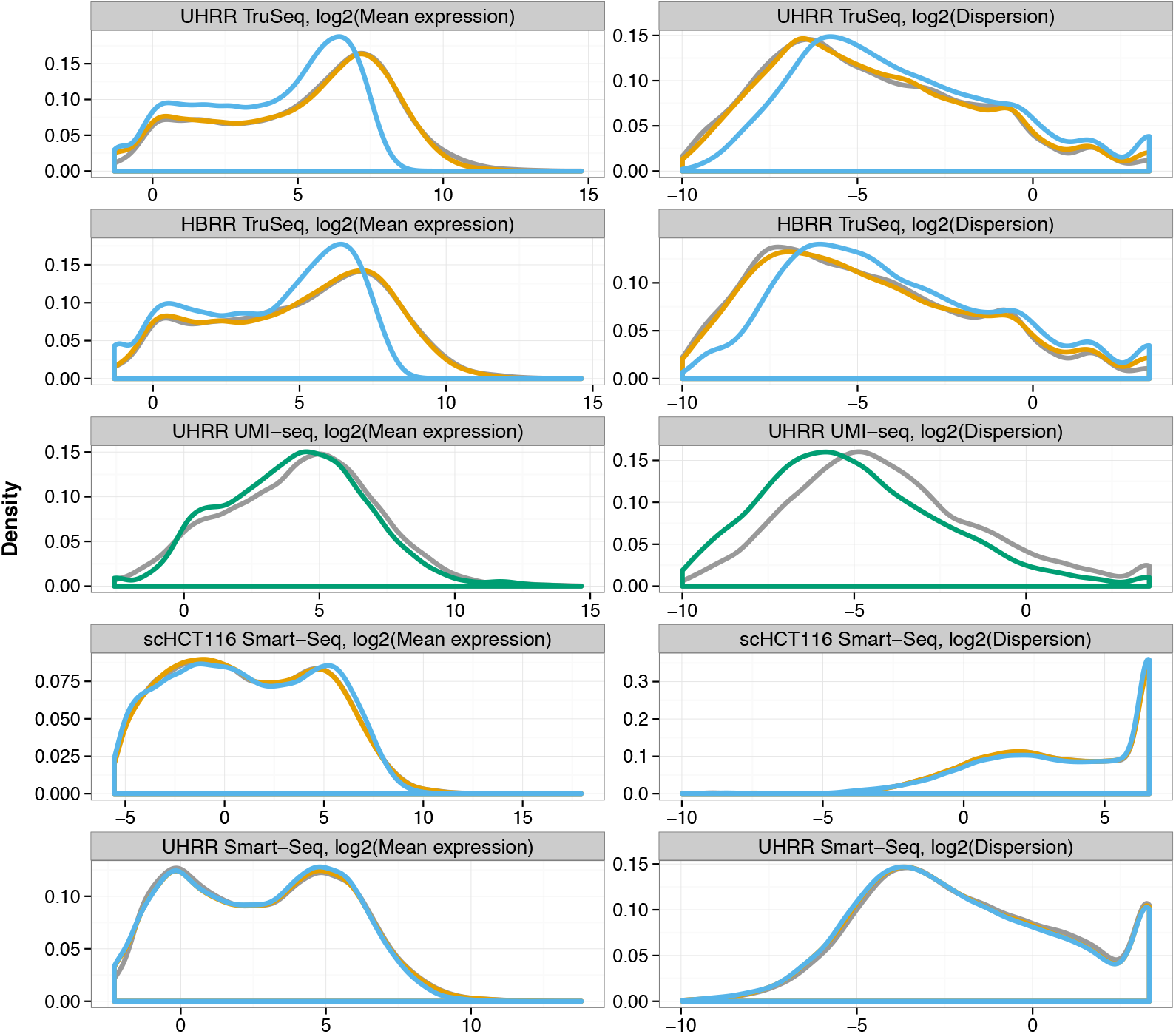
Empirical mean and dispersion distributions are used to estimate power to detect differential expression. The left panel shows density plot of log2(mean baseline expression) and the right panel the log2(dispersion) measured by DESeq2 for each study. Different duplicates treatments are represented by colors, All reads- grey, removing PE-duplicates- orange, removing SE-duplicates- blue and removing duplicate molecules in UMI-seq as green.

**Figure S3:**
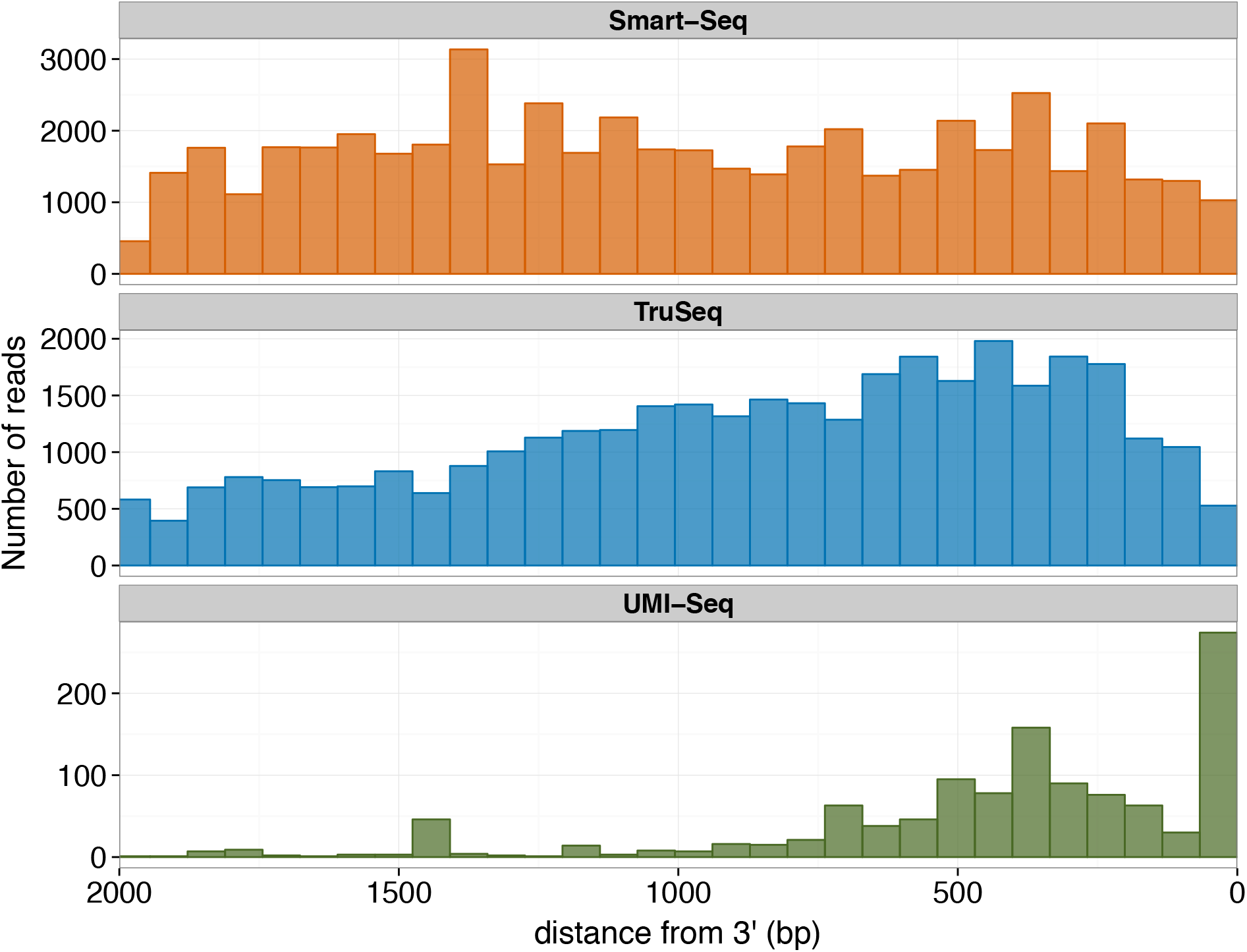
3’ bias in fragmentation site is prominent in UMI-seq. The histogram showing distance of the fragmentation site from 3’ end of the gene measured from ERCC spike-ins of length 2kb. Colors represent library preparation methods, ‘blue – Smart-Seq, ‘orange’ – TruSeq.

**Figure S4:**
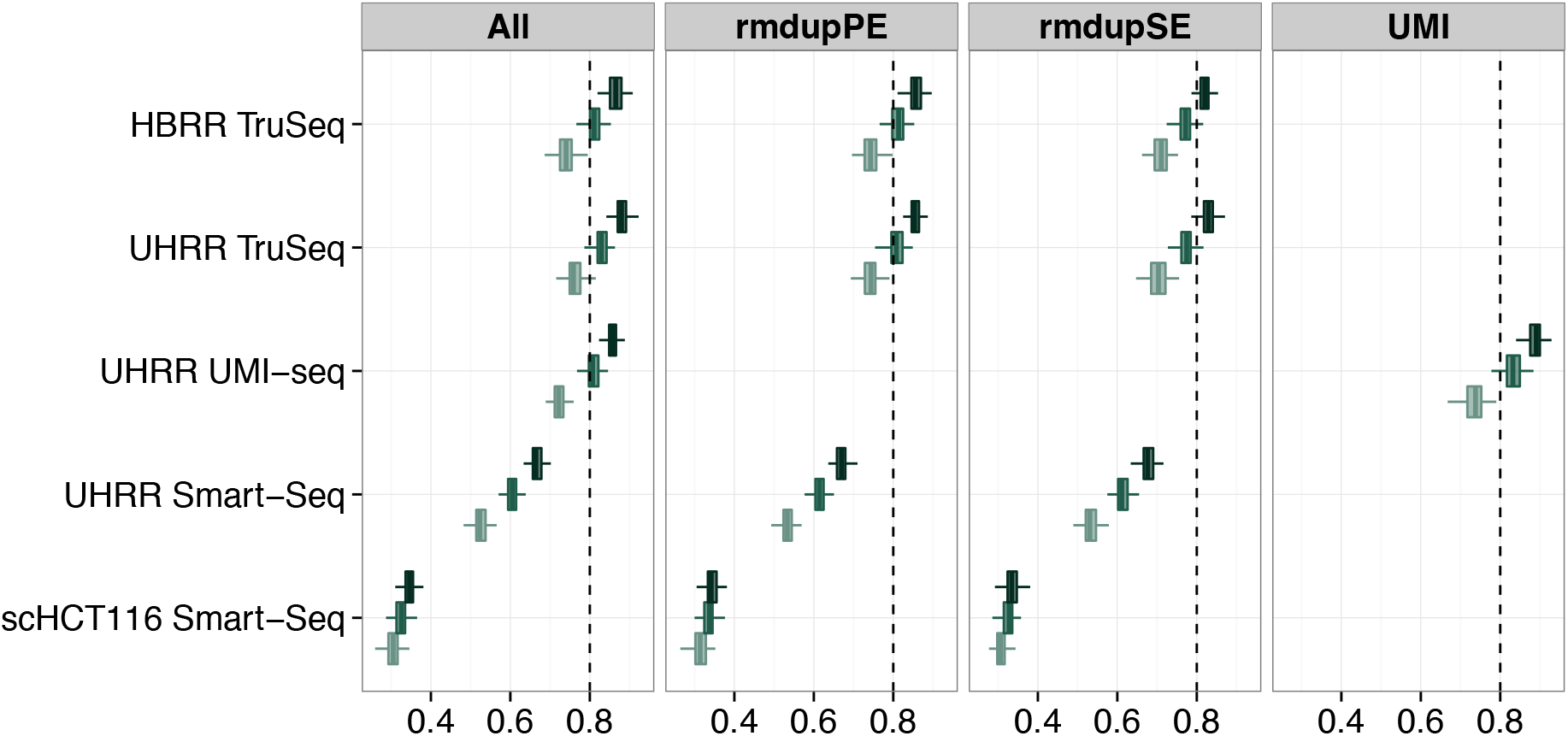
Power to detect differential expression increases with increased sample size. The box-plot shows marginal power to detect 0.5 log2foldchange at 5% nominal FDR for different sample sizes. Colors gradient from light to dark represent sample sizes 3,6 and 12 for bulk and 30,45 and 90 for single cell datasets.

## Supplementary tables

**Table S1:** Summary of squared terms from quadratic fit between PE-dup and SE-dup (PE-dup SE-dup+(SE- dup)^2^+0)

| Study name | Beta<sup>2</sup> | Std. Error | t value | Pr(> \|t\|) |
| --- | --- | --- | --- | --- |
| scHCT116 Smart-Seq | 0.6146 | 0.0309 | 19.92 | 0.0000 |
| UHRR Smart-Seq | -0.2044 | 0.6464 | -0.32 | 0.7600 |
| UHRR TruSeq | 1.0012 | 0.6212 | 1.61 | 0.2054 |
| HBRR TruSeq | 1.1487 | 0.2229 | 5.15 | 0.0142 |

**Table S2:** Median *R*^2^ and percentage of significant genes for the lm fit between GC content/Tn5 motif score and start coverage

| Study name | GC |  | Tn5 |  |
| --- | --- | --- | --- | --- |
|  | R<sup>2</sup> | %Significant* | R<sup>2</sup> | %Significant* |
| scHCT116 Smart-Seq | -0.00027 | 16% | 0.00112 | 49% |
| UHRR Smart-Seq | 0.00020 | 19% | 0.00174 | 59% |
| UHRR TruSeq | 0.00614 | 57% | 0.00077 | 43% |
| HBRR TruSeq | 0.00657 | 61% | 0.00077 | 43% |

\*Percentage of genes with p-value $\leq 0.05$

**Table S3.**
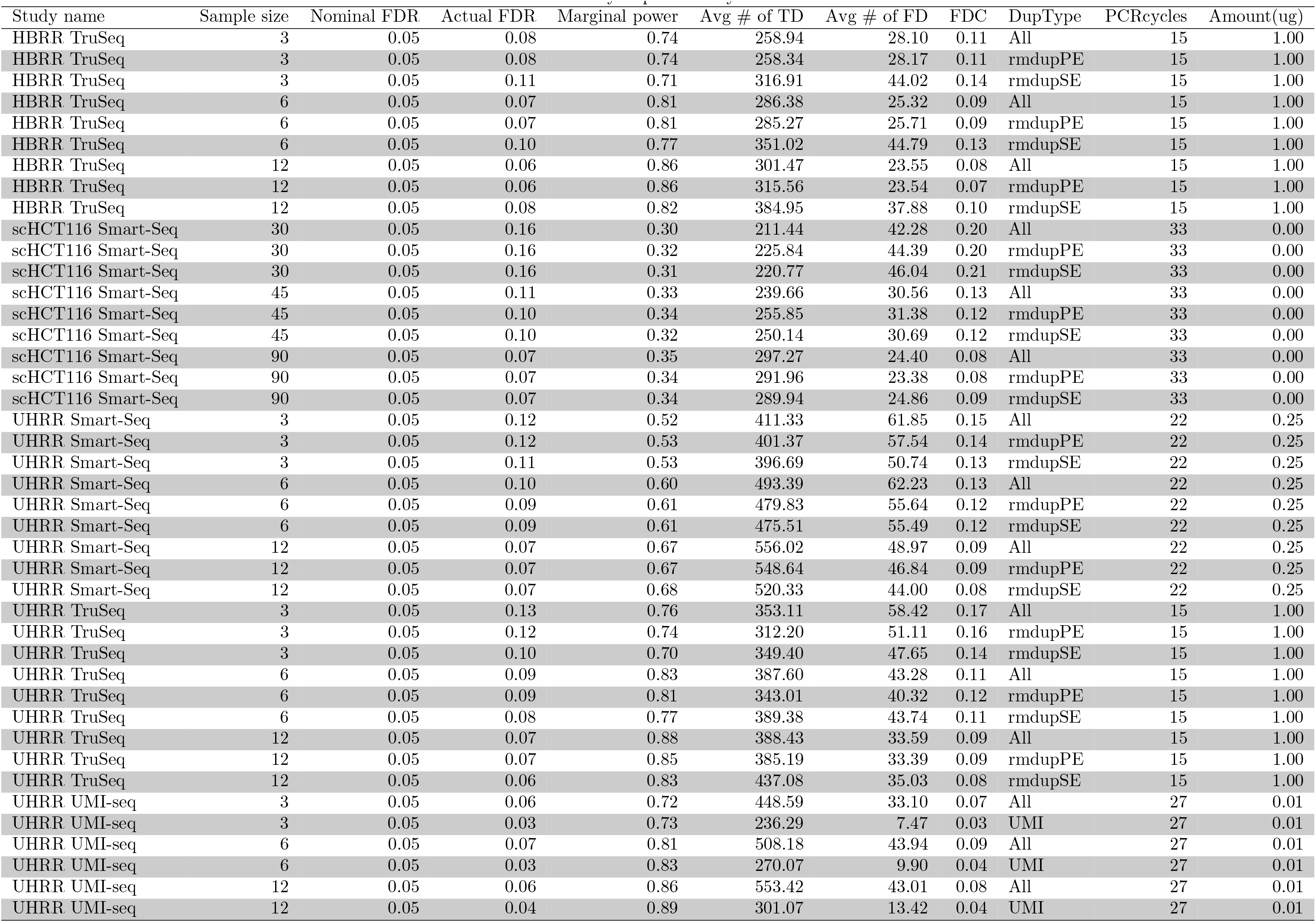
Summary of power analysis

